# Receptor-independent membrane mediated pathways of serotonin action

**DOI:** 10.1101/2020.07.01.177451

**Authors:** Simli Dey, Dayana Surendran, Oskar Enberg, Ankur Gupta, Sashaina E. Fanibunda, Anirban Das, Barun Kumar Maity, Arpan Dey, Mamata Kallianpur, Holger Scheidt, Gilbert Walker, Vidita A. Vaidya, Daniel Huster, Sudipta Maiti

## Abstract

Serotonin is a neurotransmitter as well as a somatic signaling molecule, and the serotonergic system is a major target for psychotropic drugs. Serotonin, together with a few related neurotransmitters, has recently been found to exhibit an unexpectedly high lipid membrane affinity^1–3^. It has been conjectured that extrasynaptic serotonin can diffuse in the lipid membrane to efficiently reach remote receptors (and receptors with buried ligand-binding sites)^4^, providing a mechanism for the diffuse ‘volume’ neurotransmission that serotonin is capable of^5–10^. Here we show that membrane binding by serotonin can directly modulate membrane properties and cellular function, independent of its receptor-mediated actions. Atomic force microscopy shows that serotonin binding makes artificial lipid bilayers softer. It induces nucleation of liquid disordered domains inside the raft-like liquid-ordered domains in a ternary bilayer displaying phase separation. Solid-state NMR spectroscopy corroborates this data, revealing a rather homogeneous decrease in the order parameter of the lipid chains in the presence of serotonin. In the RN46A immortalized serotonergic neuronal cell line, extracellular serotonin enhances transferrin receptor endocytosis, an action exerted even in the presence of both broad-spectrum serotonin receptor and transporter inhibitors. Similarly, it increases the binding and internalization of Islet Amyloid Polypeptide (IAPP) oligomers, suggesting a connection between serotonin, which is co-secreted with IAPP by pancreatic beta cells, and the cellular effects of IAPP. Our results uncover a hitherto unknown serotonin-bilayer interaction that can potentiate key cellular processes in a receptor-independent fashion. Therefore, some pathways of serotonergic action may escape potent pharmaceutical agents designed for serotonin transporters or receptors. Conversely, bio-orthogonal serotonin-mimetics may provide a new class of cell-membrane modulators.

## Main

The neurotransmitter serotonin, in addition to its directed action through synaptic signaling, has an indirect neuro-modulatory role^11–16^. A significant amount of serotonin is released from extrasynaptic areas, including from the cell body, away from the site of post-synaptic receptor densities^6, 7, 17–19^. Even when they are released at the synapses, a large amount of the freed serotonin diffuses away from the synaptic cleft^20^. From our current understanding, it appears that this temporally and spatially diffuse “volume neurotransmission”^21, 22^ is a wasteful attempt to reach far-away targets. However, a recent discovery of the high affinity of serotonin for lipid bilayers^1^ suggests that the lipid membrane may be the ubiquitously present target for such release. Indeed, it has been speculated that attachment and two-dimensional diffusion of serotonin in the lipid membrane may facilitate distal receptor binding^23^. Here, we hypothesize and demonstrate that passive serotonin-membrane interaction can give rise to an entirely receptor-independent pathway for modulating specific cell functions.

The serotonin-induced changes in the mechanical properties of the membrane suggest a mechanism by which it can, in principle, affect cellular function. Serotonin is present at a very high concentration (about 300 mM) in the synaptic and somatic vesicles^24, 25^. Therefore, both the vesicular and the plasma membranes (near the release sites) are exposed to a considerable amount of serotonin. Small amphipathic molecules, for example, specific anesthetics, are known to alter the physical properties of the membrane, such as membrane fluidity^26–30^. If serotonin also does so, then that in turn can affect cellular functions, including membrane protein function, membrane affinity of other molecules as a prerequisite for subsequent receptor binding, and key cellular processes such as exocytosis and endocytosis.

Here we used a broad array of biophysical tools to probe whether the binding of serotonin changes membrane mechanical properties and the degree of local molecular order. We then examine the effect of such modulation on membrane binding by proteins known to be co-secreted with serotonin, and also the rate of constitutive endocytosis, to understand the effect that serotonin can have independent of serotonergic receptors. We show that passive binding of serotonin to the membrane does indeed modulate cellular function, uncovering a novel manner in which this neurotransmitter can also influence biological function.

## Results

### 1. Effect of serotonin on mono- and biphasic lipid bilayers

#### 1.1. Serotonin interacts with model lipid membranes

We measure the binding of serotonin to small unilamellar lipid vesicles (SUVs) of two different lipid compositions: a zwitterionic mix of DOPC (1,2-dioleoyl-*sn*-glycero-3-phosphocholine): Egg Sphingomyelin: Cholesterol=2:2:1 (DEC221), and a negatively charged mix of POPC (1-palmitoyl-2-oleoyl-*sn*-glycero-3-phosphocholine): POPG (1-palmitoyl-2-oleoyl-*sn*-glycero-3-[phospho-rac-(1-glycerol) : Cholesterol=1:1:1 (PPC111). We note that membrane electrostatics appears to be important for serotonin binding^1, 31^. We test the degree of membrane-binding of serotonin using a dialysis retention assay described earlier^32^. Both SUV dispersions of PPC111 and DEC221 were incubated with 4.1 mM serotonin for 1 h, and then dialyzed against Mili-Q water using a 100 kDa MWCO membrane permeable for serotonin (but not for the SUVs) for 18 hrs. Vesicle-bound serotonin would not be allowed to diffuse out of the dialysis tubing. A control sample contains only 4.1 mM serotonin (no SUVs). Fluorescence spectra (excitation 270 nm, emission 300 −500 nm) of the dialyzed solution yield the concentration of the remaining serotonin in the dialysis tube. Excess serotonin fluorescence compared to the control sample, provides the amount of membrane-attached serotonin (Fig. S1A). The concentration of serotonin remaining after dialysis is determined from a calibration curve obtained from known concentrations of serotonin (Fig. S1B). We observe that 0.24% of serotonin remains in the DEC221 solution, 3.53% is retained in the PPC111 solution, and only trace amount (^~^0.005%) is retained in the solution without vesicles. This implies that serotonin binds strongly to the lipid membrane, and binding was dependent on the lipid charge. Assuming no lipid losses during the whole process, the partition coefficient of serotonin is calculated to be 90 and 1500 in DEC221 and PPC111, respectively. This is in reasonable agreement with the reported values^1^.

We used fluorescence microscopy to verify whether serotonin also binds to supported lipid bilayers (SLB). We stain serotonin with the fluorogenic label OPA (*ortho*-phthalaldehyde), whose effectiveness in detecting serotonin is demonstrated recently^33^. At room temperature, the DEC221 bilayer forms two different coexisting phases^34^. The phase enriched with sphingomyelin and cholesterol is more ordered and is known as the Liquid Ordered (L_o_) phase^35^. The other phase is more disordered and is dominated by DOPC known as the Liquid Disordered (L_d_) phase. A 100 μM serotonin solution is incubated with biphasic SLBs of DEC 221, which are characterized by the co-existence of L_o_ and L_d_ domains for 30 minutes and then washed with water to remove unbound serotonin. 100 μM of OPA is then added to this SLB and incubated further for 30 mins. The sample is imaged in a confocal microscope (excitation at 488 nm, and collection from 500 to 650 nm, see supplementary Fig. S2B). The fluorescence image of a control solution, identical in all respects except that serotonin is absent, is shown in supplementary Fig. S2A. The image clearly shows that serotonin strongly binds to the SLB. It is apparent from the image that serotonin preferably binds to the disordered domain (also confirmed by simultaneous AFM-confocal microscopy measurement) compared to the ordered domain (Fig. S2 D, E). The ratio of serotonin bound to the disordered vs. the ordered domain is approximately 4 (supplementary Fig. S2C).

#### 1.2. Serotonin binding reduces membrane stiffness

We perform atomic force microscopy on the SLBs to probe whether serotonin binding has an effect on the mechanical properties of the membrane. We measure membrane stiffness by determining the force of indentation, i.e., the force needed to rupture the bilayer (a representative force curve is shown in Fig. 1A. with an arrow marking the force of indentation). This force of indentation of a bilayer is first determined in water in the absence of serotonin, and then after incubating the same bilayer with 4.1 mM serotonin for 60 min. For the monophasic PPC111 bilayer, the results are presented as a histogram (Fig. 1B, n = 1410 force curves), and the average force is shown in Fig. 1C. Serotonin reduces the indentation force by 58 ± 19%. As a control experiment, we perform similar measurements in the absence and in the presence of 4.1 mM glutamate, an excitatory neurotransmitter (supplementary Fig. S3). In contrast to the effect of serotonin, glutamate does not cause any observable alterations in membrane stiffness. The pH inside an intracellular neurotransmitter vesicle is about 5.5, while that on the plasma membrane is 7.4. So we also test whether serotonin can change the pH of the solution and whether just a change of pH could affect the force of indentation. We found these effects to be minimal, as described in supporting Figs. S4 and S5.

**Fig. 1(A):**
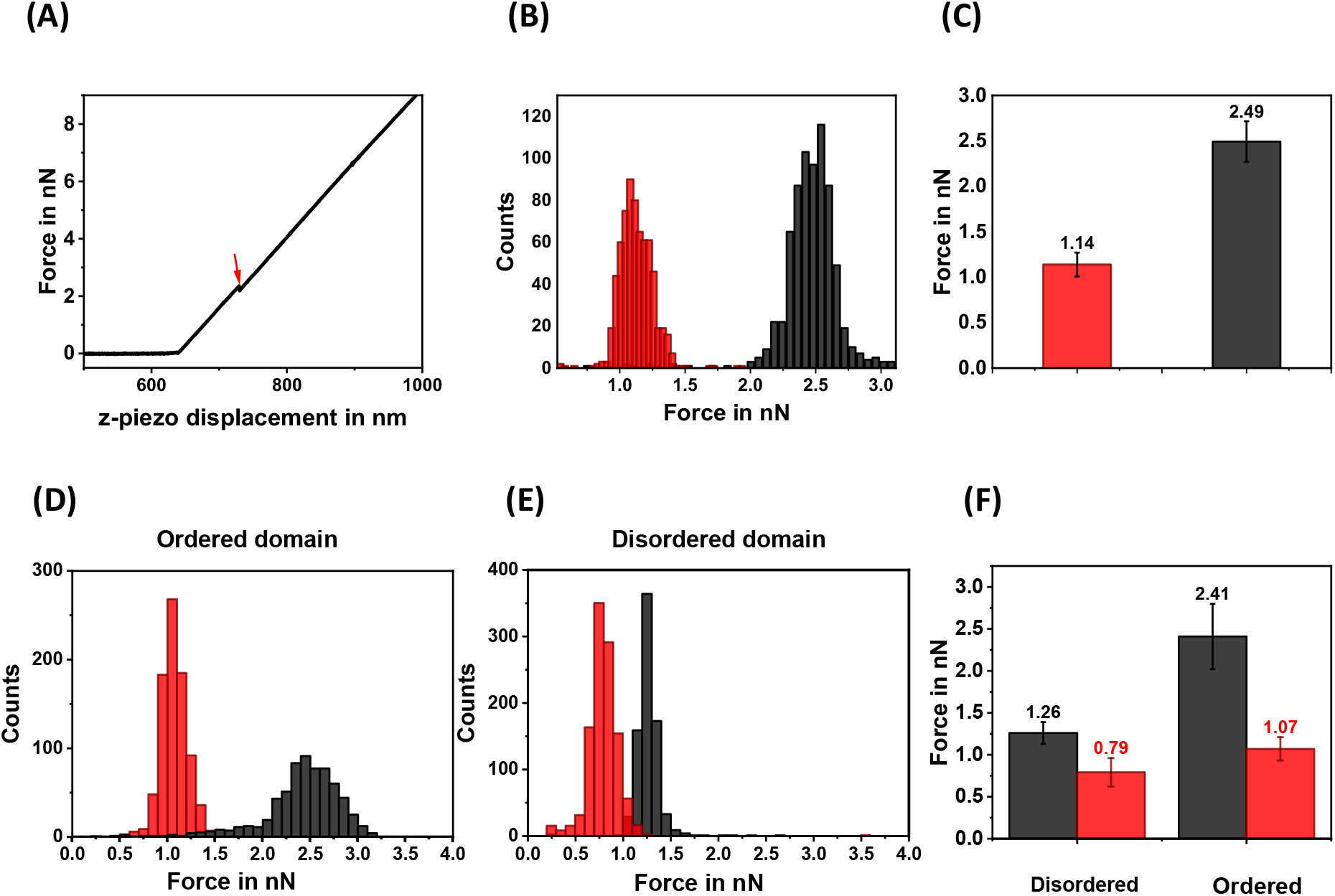
A typical indentation force curve obtained from supported POPC/POPG/Chol (1:1:1) lipid bilayer on mica. The force applied by the cantilever is plotted as a function of the z-piezo displacement. The discontinuity in the force curve marks the indentation force (red arrow). (B) Representative histogram of indentation forces before (black) and after (red) serotonin incubation on a negatively charged (PPC111) bilayer. (C) Average values of force histograms. (D), (E) representative histograms of indentation forces measured on the ordered and disordered domains respectively on a neutral biphasic (DEC221) bilayer. (F) Average values of force histograms before (black) and after (red) serotonin incubation.

Local membrane order can be an important parameter determining cell signaling, and so we want to probe the effect of serotonin on phase-separated lipid bilayers. AFM imaging characterizes the co-existence of the two phases in the DEC221 bilayer (as described in methods). We obtain ordered lipid domains with diameters on the order of a few μm. The height difference between L_o_ and L_d_ matches well with the previously reported values (from ^~^0.8 nm to 1.0 nm^36^). For indentation measurements on this biphasic bilayer, we first map the individual L_o_ and L_d_ domains and then collect traces from the AFM images. The resulting histograms are shown in Fig. 1D (for ordered domains) and Fig. 1E (disordered domains). The resulting force distribution is shown in Fig. 1F. We see that in the absence of serotonin, the L_o_ domains in the bilayer are stiffer (i.e. the indentation force is higher) compared to the L_d_ domains, as is expected. In the presence of serotonin, the force of indentation decreases in both the domains, but the decrease is much higher for the ordered domains. The force in the L_o_ domain decreases by 52.0 ± 8.3 % while that in the L_d_ domain decreases by 32.0 ± 10.3% (n = 5508 force curves from 2 different bilayers).

#### 1.3. Serotonin shrinks ordered domains in phase-separated bilayers

Biological cell membranes display a transient domain structure that is believed to play a major role in protein trafficking, in the interaction of the membrane with the cytoskeleton, and in cell signalling^37,38^. Our observation that the indentation force of the L_o_ domains of DEC221 decreases drastically may suggest that serotonin can have an effect on the relative stability of the ordered and disordered domains. We characterized the area and the perimeter of the L_o_ and L_d_ domains before and after incubation with serotonin. We observe that serotonin induces nucleation of disordered domains within the ordered domains (Fig. 2, see arrows showing a few representative sites). There is an overall decrease of the area of the ordered domain. A control sample shows no such changes over the same amount of time (supplementary Fig. S6). Also, we obtain a measure of the surface tension at the domain interface by dividing the surface area of individual domains by their perimeters. We find that the surface tension decreases by 34.0 ± 2.1% in the presence of serotonin (n = 133 L_O_ domains). This is consistent with the decrease of the force of indentation found earlier.

**Fig. 2:**
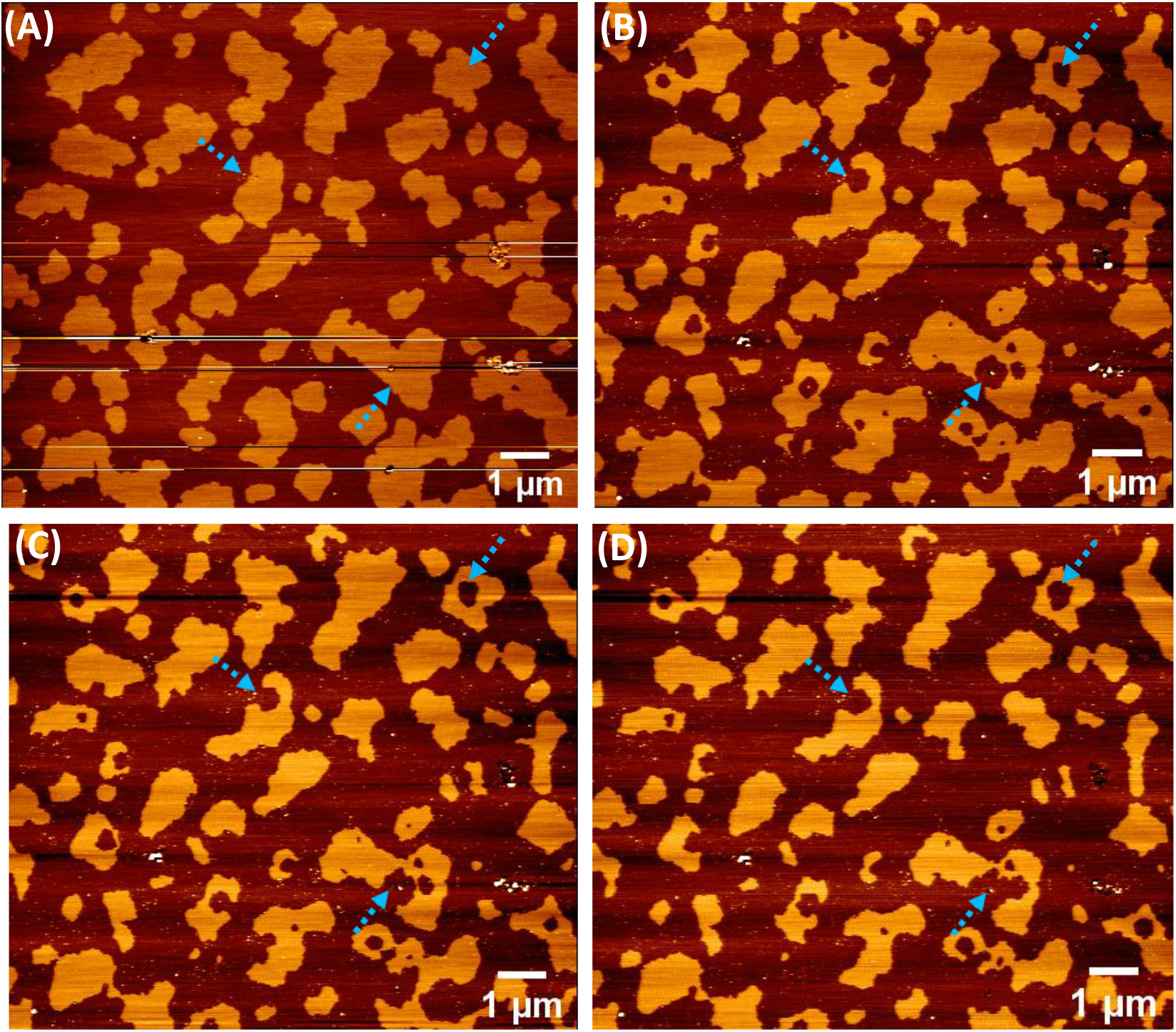
AFM images displaying time-dependent nucleation of disordered domains within the ordered domains in a phase separated lipid bilayer (DEC221) (A) 0 min (B) 20 min, (C) 120 min, and (D) 300 min after incubation of 4.1 mM of serotonin.

#### 1.4. Serotonin promotes lipid chain disorder

Solid-state NMR can probe the molecular order along lipid chains. We perform solid-state NMR experiments on multilamellar PPC111 vesicles containing serotonin and determine the order parameters of the lipid protons. We record the ^2^H NMR spectra of either chain deuterated POPC-*d*_31_ or POPG-*d*_31_ in the mixture in the presence of 0, 10, and 25 mol% serotonin (supplementary Fig. S7). From the NMR spectra, order parameter plots for each deuterated lipid at 25 and 37°C are determined and are shown in supplementary Fig. S8. We observe that in the presence of serotonin, lipid chain order is homogeneously decreased along the entire *sn*-1 chain of both POPC and POPG. Overall, the presence of serotonin decreases the average chain order parameter by 3 to 19%. At 25°C, 10 mol% serotonin does not alter the order parameter profile of POPC-*d*_31_ significantly. However, a drastic decrease in chain order is observed in the presence of 25 mol% serotonin. At 37°C, the chain order decrease is identical for both serotonin concentrations. POPG-*d_31_* responds more drastically to the presence of serotonin at both temperatures, but only minor differences are observed between the two serotonin concentrations probed here (shown in Fig. 3A). The decrease in lipid chain order leads to a decrease in the average chain length of the lipids. These serotonin-induced chain length alterations can be precisely calculated for both phospholipids of the mixture using the mean torque model^39^. The average chain lengths of both POPC and POPG of the mixture in the absence and in the presence of serotonin is plotted in Fig. 3B. Serotonin causes the lipid chains to decrease in length by ΔL, where ΔL is between 0.3 and 0.9 Å. This decrease in the lipid chain length is due to a serotonin-induced increase in the number of *gauche* conformers in the chains.

**Fig. 3(A):**
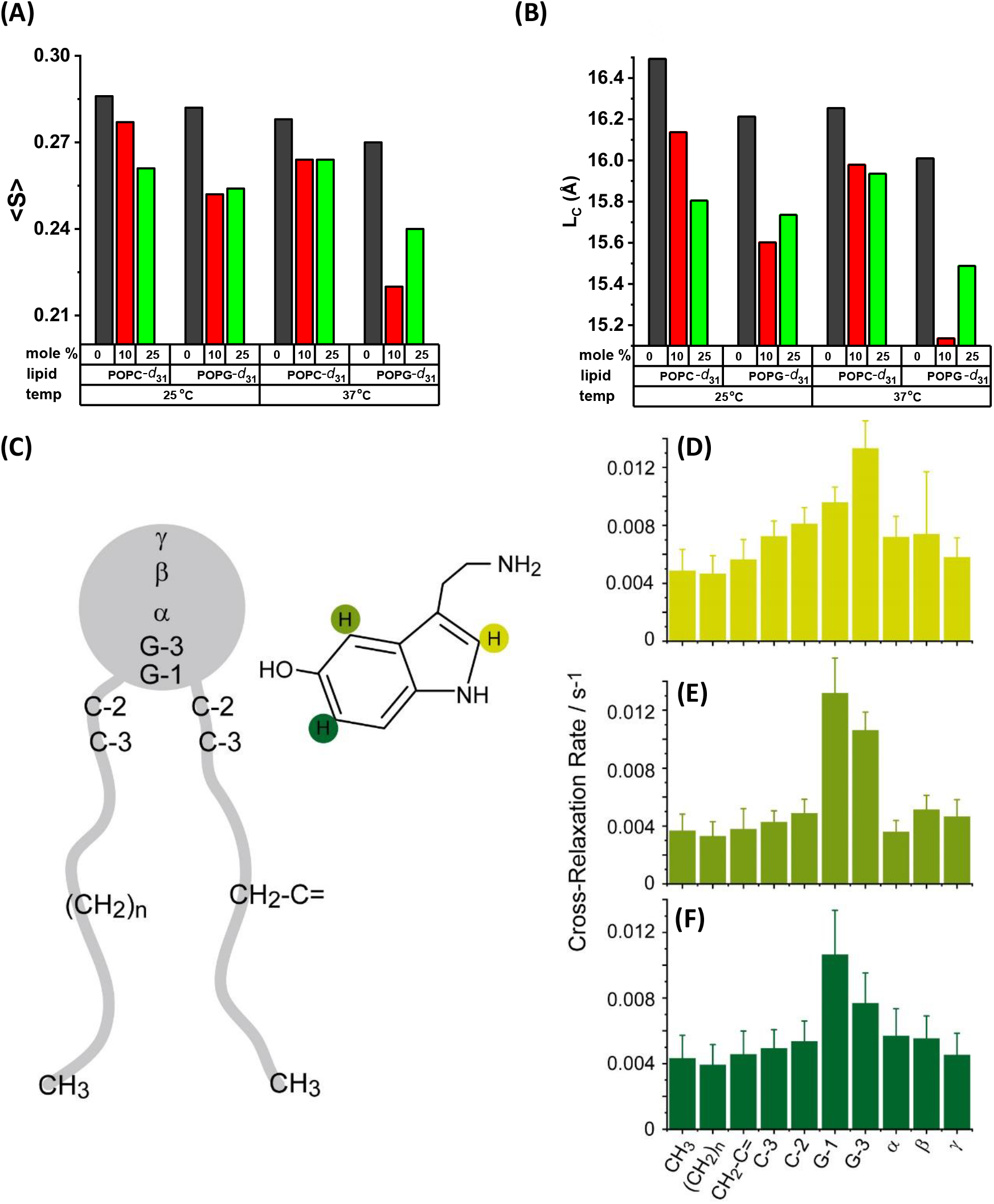
^2^H NMR average order parameters <S> of POPC-d_31_:POPG:Chol(1:1:1) and POPC:POPG-d_31_:Chol(1:1:1) with 0, 10, 25 mol% of serotonin at (i) 25°C and (ii) 37°C. (B) average lipid chain length of the above mentioned membrane composition. (C) schematic representation of serotonin interaction with POPC lipid chain segment. (D,E,F) are the ^1^H NOESY NMR cross-relaxation rates representing the contact probability between the individual protons of serotonin labeled in (C) with the respective lipid segments.

We then probe the average location of serotonin in POPC membranes using ^1^H NOESY NMR. This technique is well-suited to localize a small lipophilic molecule in the membrane and to determine its distribution parallel to the membrane normal^40^. The cross-relaxation rates represent the contact probability between the individual protons of serotonin with the respective lipid segments. Our results show that the ring system of serotonin is broadly distributed within the membrane with a maximum found in the glycerol region, which is at the lipid-water interface of the membrane (Fig. 3 C-F). The molecular basis of the disordering effect of serotonin on lipid acyl chains can be understood from these results. Serotonin inserts into the lipid membrane and intercalates between neighboring lipid molecules in the glycerol region. This creates free volume in the acyl chain region of the membrane, which is occupied by larger amplitude motions of the chains, resulting in the observed lowering of the chain order parameters. The NMR results corroborate the decrease in the stiffness of the membranes observed by AFM. We note that this effect may depend on the nature of the lipid, since serotonin may distribute differently in bilayers of different compositions.

### 2. Effect of serotonin on cell membranes

#### 2.1. Serotonin increases membrane binding of amyloid oligomers

We hypothesize that the observed alterations in the mechanical properties of the membrane may contribute to the action of serotonin on cells. An increase of membrane disorder may facilitate the binding of small extracellular molecules and peptides to the membrane, and this may have physiological consequences. Furthermore, membrane protein function is strongly related to the elastic properties of the bilayer^41^. We probed the affinity of IAPP oligomers (synthesis of which is described previously^42^) to the membrane of live cells in the presence of extracellular serotonin. IAPP is an amyloidogenic peptide that is co-secreted with serotonin from pancreatic beta cells, and its membrane interaction has been linked to Type II diabetes^43–45^. We have already shown that the oligomeric form of this peptide has a large membrane affinity, which may partly explain its biologically toxic role^42^. IAPP oligomers prefer to bind to the disordered phase of the lipid bilayer (unpublished), which in the light of the current results suggests that IAPP binding to the cell membrane may be modulated by serotonin. However, most mammalian cells have receptors for serotonin, so serotonin can, in principle, elicit a direct receptor-mediated response. Here we have used the RN46A cells, which is an immortalized serotonergic cell line derived from rat raphe nuclei^46^. With the help of polymerase chain reaction (PCR), we determine that the major serotonergic receptors present in the RN46A cells are the serotonin 5-HT_1A_ and 5-HT_2C_ (supplementary Fig. S9). mRNA expression levels of other 5-HT receptors (5-HT1B, 5-HT3, 5-HT4, and 5-HT7 receptors) are not significant. To block 5-HT_1A_ and 5-HT_2C_ receptors, as well as the serotonin transporters (SERTs), we pre-treat the cells with WAY100635^47^ (10 μM), Methysergide (10 μM), Ketanserin^48^ (10 μM) and Fluoxetine (10 μM) in all cell experiments. The cells are then washed with fresh Thomson’s buffer (TB) and incubated with 200 nM of rhodamine-labeled IAPP oligomers for 30 minutes. The cells are washed again with fresh TB and imaged in TB using a confocal microscope (Zeiss 880, Germany). The average fluorescence intensity of the cell region is analyzed after subtraction of the non-cell background, using ImageJ software^49^, and reports the extent of IAPP binding. The results are depicted in Fig. 4A and 4B. We see that IAPP binding increases by 32.0 ± 10.6% (p < 0.05) when the cells are incubated with 0.5 mM serotonin. We use a 10× lower concentration of serotonin compared to the bilayer experiments, as we do not want to cause the large changes of membrane stiffness observed in those studies. Cells treated with just the blockers do not exhibit any change in intensity (supplementary Fig. S10C). We note that serotonin also quenches rhodamine fluorescence. Therefore, the actual increase, which is measured by the increase of fluorescence at the membrane and inside the cell, is likely larger than the measured increase. Supporting Fig. S11 describes a separate experiment which characterizes the quenching of PPC111 lipid membrane-bound Rh-IAPP by serotonin. The extent of quenching at the concentration level used in these biological experiments is 49%. So we estimate that the actual level of increase of IAPP binding is 2.6 ± 0.3 times. Collectively these experiments show that serotonin binding can strongly alter the affinity of cell membranes to extracellular proteins and peptides.

**Fig. 4:**
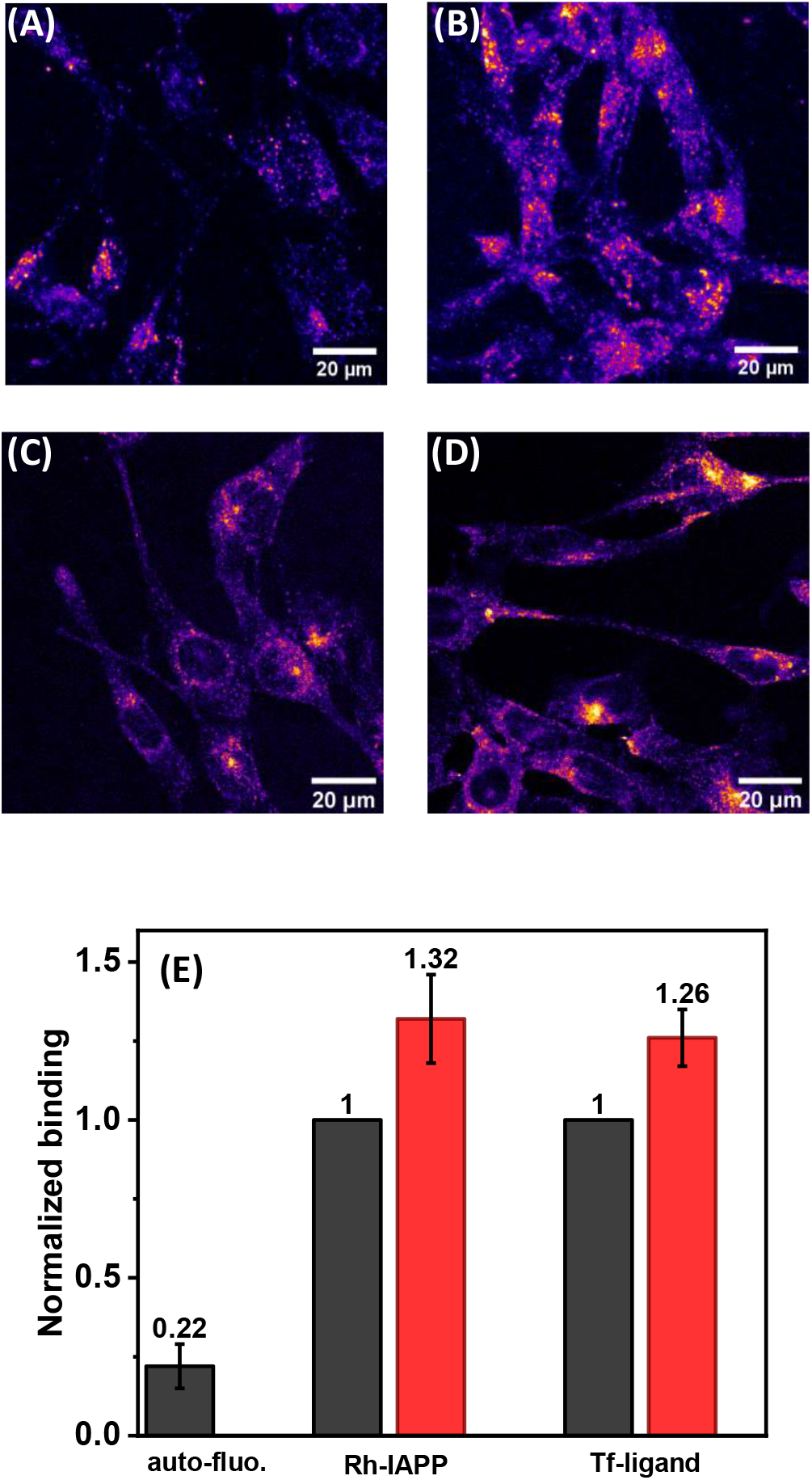
Effect of serotonin on the interaction of RN46A cells with IAPP (IAPP) and transferrin. (A,B) Confocal images of the binding and internalization of Rhodamine (Rh)-labelled IAPP oligomers to RN46A cells in the absence (A) and in the presence(B) of serotonin. (C,D) Confocal images of transferrin receptor internalization by RN46A cells showing AlexaFluor488-transferrin distribution in absence (E) and in presence (F) of serotonin. (E) analysis of the extent of binding of different probes in absence (black) and in presence (red) of serotonin. Cells are pre-treated with 5-HT_1A_ receptor antagonist, WAY100635 (10 μM), 5-HT2 receptor antagonists Methysergide (10 μM) and Ketanserin (10 μM), and SERT inhibitor Fluoxetine (10 μM).

#### 2.2. Serotonin enhances endocytosis

We also probe whether serotonin modulation of the membrane properties affects the process of endocytosis. Endocytosis requires the generation of large membrane curvatures, which should be gacilitated by the decrease of the membrane stiffnes. So one would predict that the rate of endocytosis may be enhanced by seeotonin. We use a constitutive endocytosis process, not known to be related to serotonin signalling, as a measure for the modulation of endocytosis rates. We follow the rate of endocytosis of the transferrin (Tf) receptor using a fluorescent transferrin conjugate (Alexa Fluor488-transferrin). The experiments are similar to those described for IAPP binding and are also performed in the presence of the serotonin transporter inhibitor (fluoxetine) and receptor antagonists (Way100635, Methysergide, and Ketanserin). The integrated brightness of the cell after 15 minutes of incubation with Alxa488-Tf is used as a measure of the rate of endocytosis. The images are recorded with a confocal microscope using 488 nm Argon laser excitation. The total intracellular intensity is calculated by z-projecting the image stacks and subsequently delineating the cell boundaries manually. We observe that the level of transferrin internalization goes up by a factor of 26.0% ± 7.1% (p < 0.05) in presence of serotonin, as shown in Fig. 4C and 4D. This indicates that serotonin-membrane interaction can affect membrane proteins unrelated to serotonin, potentially providing a membrane-mediated pathway for modulating the physiological states of cells.

## Discussion

Serotonin is water-soluble to >2 M concentration, yet it attaches to lipid membranes with a partition coefficient between 90 and 1500. Vesicular membranes and plasma membranes near serotonin release sites are expected to contain a considerable concentration of serotonin. Here we address whether serotonin affects the properties of the membrane. This question assumes significance because any change of membrane properties will likely affect fundamental cellular processes that are mediated by the membrane, such as endo- and exocytosis, and dynamics of membrane proteins.

High membrane partitioning of serotonin is observed in DMPC and DOPC vesicles^1^. Here we probe whether this high affinity is also observed for multicomponent membranes containing neutral (DEC221) and negatively charged lipids (PPC111), and also containing sphingomyelin and cholesterol (Fig. S1). Our vesicle retention experiment (Fig. S1A) confirms that high membrane partitioning is also true for these two types of membrane mixtures, though to different degrees. The molar serotonin-to-lipid ratio in vesicles exposed to 4.1 mM serotonin is 0.7% for the zwitterionic lipid vesicles, while it is 11% for the negatively charged vesicles. Thus the substantial quantity of serotonin is present in the membrane, and depending on their location in the membrane, they can be expected to have significant effects on membrane properties.

Two of the major properties of a membrane are its stiffness and local molecular order. These parameters can influence the membrane traffic and also the functioning of membrane proteins, which undergo large conformational changes during their function requiring membrane elasticity, which is modulated by the lipid dynamics. We measure the influence of serotonin on both of these properties. The negatively charged PPC111 membrane shows a profound lowering of its stiffness, going from an average of 2.49 ± 0.22 nN to 1.14 ± 0.13 nN (Fig. 1C). The zwitterionic biphasic DEC221 bilayer also shows a decrease, but mostly in the stiffness of the ordered domain. Significantly, serotonin decreases the area of the ordered domains (Fig. 2), and nucleates disordered phases within the ordered ones. These cholesterol and sphingomyelin rich ordered domains are representative of raft-like structures on the cell membrane, which are important for cellular signaling processes^38, 50, 51^. Serotonin-induced reduction of the fractional area of the ordered domain can, therefore, have significant consequences for cell signaling.

Small molecules, such as cholesterol or anaesthetics^12, 34, 52^, can change membrane properties, depending on the location of the molecule in the lipid bilayer. Molecular dynamic simulations suggest that serotonin primarily localizes near the headgroup part (between the headgroup and the glycerol parts to be precise) of the membrane^1^. Such a location can promote lipid chain disorder, especially in the lower parts of the lipid chains. We experimentally investigate the location of the serotonin molecules and its effect on lipid disorder by using solid-state NMR spectroscopy. Our results show that serotonin can be distributed throughout the membrane, but its average location is close to the glycerol group of the phospholipids. The order parameter measurement reveals that the disorder of the lipid chains indeed increases considerably (Fig. 3), and the lipid chain effectively shortens by about 0.5 Å. These results are clearly in agreement with the increase of disorder observed in our AFM force measurements.

Such enhancement of disorder should have a bearing on membrane-mediated processes in a cell. However, it is not straightforward to exclusively measure the membrane-mediated effects of serotonin. Most neurons, and indeed, most cell types in the human body have serotonin receptors^53^, so the effects will be dominated by serotonin receptor-driven signaling. We determine the types of serotonin receptors present in the RN46A serotonergic neuronal cell line used for our study and carry out all our cellular measurements under adequate concentrations of antagonists for these serotonergic receptors (as well as the serotonin transporters). An increase in membrane disorder can be expected to facilitate the insertion of molecules with high membrane affinity. We have separately observed that IAPP oligomers prefer the disordered regions of a biphasic lipid bilayer by a factor of 7:1, compared to its ordered regions (Gupta and Dey, unpublished). We observe that binding and uptake of IAPP oligomers (associated with Type II diabetes) by the cells are indeed enhanced in the presence of 0.5 mM of serotonin (Fig. 4 A and 4B). It is interesting to note that serotonin is co-secreted by the pancreatic beta cells together with IAPP^45, 54–56^, so the effect of serotonin on the degree of membrane attachment of IAPP could be physiologically significant. A reduction in stiffness should promote exo- and endo-cytosis, by allowing the formation of larger curvatures in the membrane. We investigate the effect of serotonin binding on endo-cytosis by measuring the internalization of transferrin receptors. The transferrin receptors have no known interactions with serotonin, yet we observe a considerable increase in their endocytosis in the presence of serotonin (Fig. 4C, D). Thus two important membrane-mediated processes, i.e. protein binding and the rate of endocytosis, are both strongly modulated by serotonin. This indicates that critical cellular processes may respond to serotonin via membrane-mediated effects in addition to the more traditional receptor-mediated signaling pathways.

It may be asked whether the concentrations of serotonin used in our experiments (^~^4 mM for the studies on artificial bilayers, 0.5 mM for cell experiments) are physiologically relevant. The serotonergic vesicles are known to contain >250mM of serotonin^24, 25^, much more than the concentrations used here. So the vesicular membrane will be saturated with serotonin, and during exocytosis, these will provide serotonin-rich membrane patches for mediating activity on the plasma membrane. However, a substantial amount of serotonin is also directly available to the plasma membrane, at least transiently, near the site of vesicular release. Model calculations suggest that the release of a single small monoaminergic synaptic vesicle can elevate the extracellular concentration of monoamines to several μM for ^~^ms time scales^57^. However, cell soma can perform a sustained release of hundreds of vesicles of much larger size (with diameters >100 nm compared to the ^~^40 nm typical synaptic vesicles^24, 25, 58^. Given that serotonin strongly partitions in the membrane, such sustained release may have a cumulative effect on the membrane serotonin availability. Therefore, a ^~^mM serotonin concentration may not be unusually high for the plasma membrane near the release sites. So the biophysical and cellular effects observed here may also be expected to occur *in vivo*.

Overall, our experiments show that serotonin, at concentration levels relevant *in vivo*, can interact with the cell membrane, increase membrane disorder, and profoundly change the membrane modulated properties of cells. The membrane provides a ubiquitously present effector for serotonin, through which it can directly influence cellular physiology, in keeping with other receptor-independent roles of serotonin such as serotonylation of target proteins^56, 59^. This membrane-mediated pathway may be secondary to the conventional receptor-mediated pathways, but it will be rather important in situations, for example, where those receptors are inhibited by pharmaceutical agents. Its influence on membrane processes can also become an important consideration when the extracellular serotonin is maintained at an elevated level by serotonin transporter blockers. Our results also suggest that bio-orthogonal serotonin-mimetic molecules can provide useful pharmacological modulators of cellular properties. Therefore, a comprehensive understanding of serotonin biology and pharmacology needs to take this novel membrane-mediated pathway into account.

## METHODS

### Materials

The lipids 1-palmitoyl-2-oleoyl-*sn*-glycero-3-phosphocholine (16:0-18:1, PC), 1-palmitoyl-2-oleoyl-*sn*-glycero-3-[phospho-rac-(1-glycerol)] (16:0-18:1 PG), 1,2-dioleoyl-*sn*-glycero-3-phosphocholine (DOPC, 18:1), N-hexadecanoyl-D-erythro-sphingosylphosphorylcholine (egg SM, 16:0) as well as the in the *sn*-1 chain per-deuterated versions POPC-*d*_31_ and POPG-*d*_31_ were purchased as powder from Avanti Polar Lipids (Alabaster, AL), Cholesterol (Chol) and *ortho*-phthalaldehyde (OPA), Way100635 maleate, Methysergide maleate and Ketanserin tartrate were purchased from Sigma (St. Louis, MO). Serotonin-hydrochloride was purchased from Merck (Darmstadt, Germany). Chloroform and methanol AR graded were purchased from S. D. Fine-Chemicals Ltd. (India), and Milli-Q water (Millipore, Merck), deionized to a resistivity of 18.2 MΩ·cm^−1^, were used for all experiments. Lipid extruding kit and Nucleopore Track-etched polycarbonate membrane of 50 nm pore diameter were bought from Avanti Polar Lipids (Alabaster, AL). Biotech CE Tubing: 100 kDa MWCO dialysis membrane was purchased from Spectrum Laboratories, Inc. (MA, USA). Fluoxetine hydrochloride was purchased from Tocris Bioscience (Minneapolis, MN). Alexa Fluor488-transferrin was purchased from Molecular Probes (Eugene, OR).

### Preparation of planar supported lipid bilayers

Planar supported lipid bilayers were prepared by vesicle fusion method^60^. To prepare POPC/POPG/Chol in 1/1/1 molar ratio (PPC111), the required amount of lipids was dissolved in chloroform. For DOPC/Egg SM/Chol in 2/2/1 molar ratio (DEC221), the required amount of lipids was dissolved in 1:1 (by volume) methanol: chloroform. The solvent was evaporated under extra pure Argon flux and then subjected to vacuum for 24 hours for complete removal of organic solvents. The lipid films were rehydrated in water to a final concentration of 2.5 mg/ml. The lipid suspension was then vortexed vigorously and extruded using 50 nm pore diameter polycarbonate membrane at 60 °C. After extrusion, 40 μL of 100 mM CaCl_2_.2H_2_O, 50 μL of extruded lipid solution and 210 μL of Milli-Q water were deposited sequentially on freshly cleaved mica previously glued on to a glass coverslip affixed to a liquid incubated for 1 h at 60 °C in a water bath and slowly cooled down to room temperature. The samples were rinsed extensively with deionized water to remove non-fused vesicles. The presence of bilayer was confirmed using AFM (Atomic Force Microscopy). AFM contact mode imaging shows that the ternary mixture containing DOPC, Egg Sphingomyelin, and Cholesterol (2:2:1) forms a continuous biphasic bilayer with ^~^1 nm height difference between the two phases. The PPC111 bilayer has no phase separation and no image contrast. However, force indentation by AFM confirms the presence of the bilayer in the sample.

### AFM imaging and force indentation

All AFM measurements were acquired using the NanoWizard II system (JPK Instruments, Berlin, Germany) which is mounted on an Axiovert Inverted Microscope (Zeiss, Germany). AFM topographic images were obtained in Contact-mode, using silicon nitride cantilevers (Bruker, Camarillo, CA) with a spring constant of 0.01 N m^−1^. The force during imaging was kept as low as possible, and the scan rate was kept at 1 Hz. Height, vertical and lateral deflection error signals were simultaneously recorded for both trace and retrace. All the AFM images were analyzed using JPK image processing software, and the images were plane fitted with a 1^st^ order polynomial.

Force measurement was performed after the calibration of sensitivity, resonance frequencies (both in air and in water), and the spring constant, using the thermal noise method^61^. The cantilever used for all force experiments has a resonance frequency of 10-20 kHz and a spring constant of 0.03 N/m. Sensitivity and spring constant measurements were performed after each experiment. The sensitivity values before and after the experiments remained the same within errors. All the AFM experiments were performed on mica glued to coverslip glass in a liquid cell. The bilayer was hydrated until the end of the experiment. The force indentation experiments on phase-separated DEC221 bilayers were preceded by imaging of the bilayer, which located the L_o_ and L_d_ domains. The indentation forces were measured then by selecting a small area inside both the L_o_ and L_d_ domains. For the PPC111 bilayer, the total Z piezo displacement was 1.0 μm. The indenting speed both for approach and retraction was kept at 0.5 μm·s^−1^. For the DEC221 bilayer, the total Z piezo displacement was kept at 5.85 μm. The indenting speed both for approach and retraction was kept at 2.0 μm·s^−1^. In the non-contact region, the AFM tip first approaches the top surface of the bilayer during which the force remains constant. As it hit the bilayer surface at the contact point, the force increases. At some point, the tip started indenting the bilayer, which was followed by a sudden breakthrough, which appeared as a ‘kink’ in the smooth approaching force-distance curve (shown in Fig. 1A). This force is known as the indentation force. It was followed by the tip reaching the solid mica support. All the experiments were carried out at different positions on both DEC221 and PPC111 bilayer under similar conditions. The images and batches of indentation force curves were analyzed using JPK Data Processing software. Forces of indentation were extracted from each approach curve to build the histogram.

### Solid-State NMR spectroscopy

Lipids and serotonin, dissolved in organic solvents, were mixed and evaporated at 40°C in a rotary evaporator. The molar ratios of all components were POPC/POPG/Chol/Ser 1/1/1/0; 3/3/3/1 or 1/1/1/1. After evaporation of the solvent, samples were re-dissolved in cyclohexane and converted into a fluffy powder by lyophilization overnight at high vacuum. The samples were hydrated with 50%wt K2PO4 buffer (20 mM K2PO4, 100 mM NaCl, 0.1 mM EGTA pH 7.4). After hydration, samples were freeze-thawed 10 times with gentle centrifugation for equilibration and finally transferred to 5 mm glass vials.

^2^H NMR measurements were acquired on a Bruker 750 NMR Avance I spectrometer at a resonance frequency of 114.9 MHz for ^2^H. The two π/2 pulses were 2.35-2.5 μs, the relaxations delay 1 s, and the delay between the pulses 30 μs. A single-channel solids probe equipped with a 5 mm solenoid coil was used. Order parameter profiles were calculated after de-Pakeing the spectra according to Huster et. al^62^.

### Cell culture

RN46A cells (a rat serotonergic neuronal cell line) were cultured in Dulbecco’s modified Eagle’s media-F12 (1:1) (Gibco, USA) media supplemented with 10% heat-inactivated fetal bovine serum (Gibco, USA), 50 units/ml Penicillin and 50 μg/ml Streptomycin (HiMedia, India) in T-25 canted neck flasks. Cultures were maintained in a humidified atmosphere at 37°C with 5% CO_2_. Media was changed every 48 hours. For confocal imaging experiments, the cells were plated on poly-L-lysine (Sigma, St. Louis, MO, 0.1 mg/ml) coated coverslips and used the following day. Cells were imaged in Thomson’s buffer (TB, consisting of 20 mM sodium HEPES, 146 mM NaCl, 5.4 mM KCl, 1.8 mM CaCl_2_, 0.8 mM MgSO_4_, 0.4 mM KH_2_PO_4_, 0.3 mM Na_2_HPO_4_, and 5 mM D-glucose; pH adjusted to 7.4).

### Quantitative Real-time Polymerase Chain Reaction (qPCR) Analysis

RNA was extracted from RN46 cells using the commercially available Trizol reagent (Invitrogen, USA). 2 μg of RNA was reverse transcribed to complementary DNA (cDNA), using a cDNA synthesis kit (PrimeScript 1st strand cDNA Synthesis Kit, Takara Bio), according to the manufacturer’s instructions. cDNA was then diluted and subjected to quantitative real-time PCR, using gene-specific primers and KAPA SYBR (KAPA Biosystems), on a Bio-Rad CFX96 real-time PCR machine. 18S ribosomal RNA was used as a housekeeping gene to normalize gene expression levels, and relative expression levels were calculated by the DDCt method, as described previously^63^.

### Confocal Microscopy

Confocal Fluorescence microscopy on both bilayers and RN46A cells was performed on LSM 880 (Zeiss, Germany).

For confocal imaging of bilayers and RN46A, 488 nm and 543 nm, 633 nm excitation light were taken from Ar ++ and He-Ne laser respectively, which were reflected by a dichroic mirror (MBS 488/543/633) and focused through a Zeiss C-Apochromat 40x, NA 1.2, water immersion objective onto the sample. The fluorescence emission was collected by the same objective and sent to a GaAsP detector.

## Notes

### Competing Interest Statement

The authors have declared no competing interest.

