## Supporting Information for "Receptor-independent membrane mediated pathways of serotonin action"

**Fig. S1**

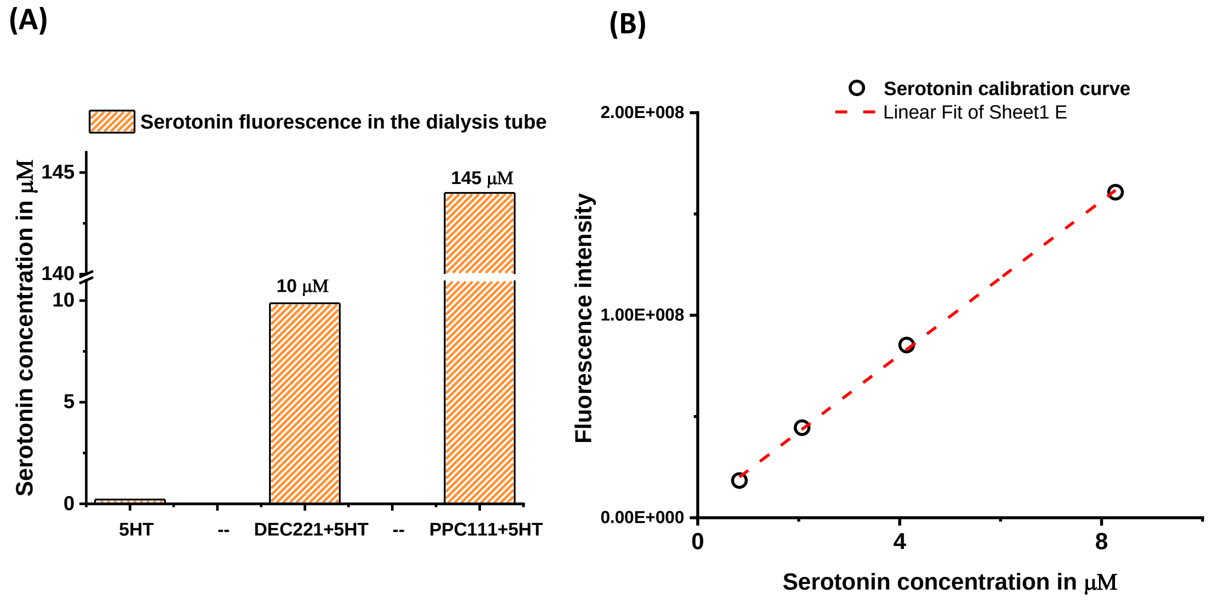


Fig. S1: Estimating the binding of serotonin with DEC221 and PPC111 vesicles by monitoring serotonin fluorescence (excitation 270 nm and emission 300-500 nm). (A) Residual fluorescence intensity of serotonin after dialysis. (B) calibration curve of serotonin obtained by plotting the fluorescence of a series of serotonin of known concentration, recorded at 340 nm.

**Calculation of serotonin partition in lipid vesicles:**

Considering no lipid loss during dialysis, the concentration of total lipid in PPC111 and DEC221 vesicles are 1304 μM and 1485 μM respectively.

Thus the partition coefficients of serotonin in PPC111 and DEC221 are ~1500 and ~90 respectively.

However, in practise, we tried to estimate the partition values from experimentally obtained quantities and we set an upper limit for these values described below. We used Fluorescence Correlation Spectroscopy to obtain such numbers. To calculate the total lipid concentration, both types of vesicles were treated with Rh-IAPP of concentration 350 nM characterised against standard Rhodamine B of concentration 125 nM.

The vesicle concentration thus obtained from FCS is 26.37 nM.

And, the total lipid per vesicle is $= \frac{Total\ surface\ area\ of\ the\ lipids\ in\ unit\ vesicle}{surface\ area\ of\ each\ lipid\ head\ group}$

The total surface area of the vesicle is 4 π ($R^2$+ $r^2$) = [where R = Outer radius (=26.43 nm, calculated considering the hydrodynamic radius of Rhodamine is 0.57 nm), r = inner radius and R-r = 4 nm i.e. the average thickness of bilayer, Avg. surface area of each lipid head group = 0.6 $nm^2$]

Hence, the estimated total lipid concentration of PPC111 vesicles =

vesicle concentration of PPC* total lipid per unit vesicle = 0.66 mM

Considering the serotonin concentration bound to PPC111 vesicles after dialysis i.e. 145 µM, the estimated partition coefficient of serotonin in PPC111 vesicles ~ 3000.

This sets the upper limit of this value which could be overestimated based on the size of the vesicle probed by IAPP (since IAPP might bind to vesicles of selective size).

The binding of IAPP to DEC221 unilamellar vesicle was rather low probed by FCS to obtain any information of serotonin partition.

**Fig. S2**

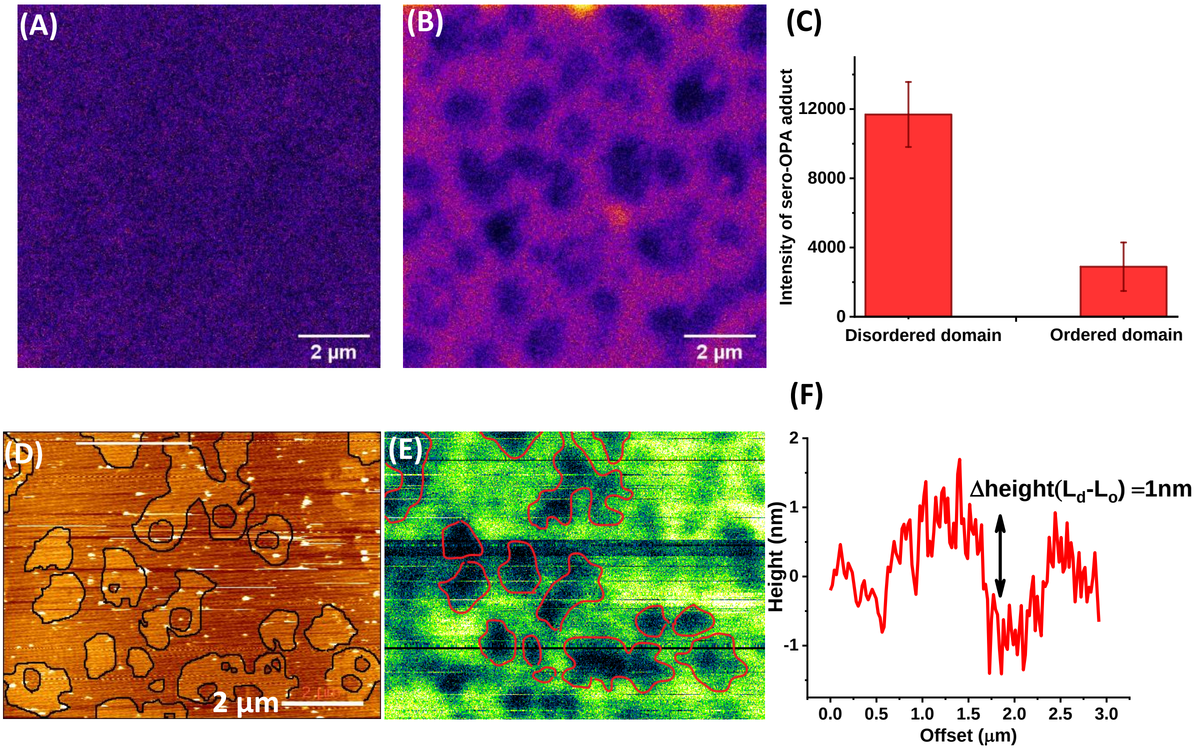


Fig. S2: Fluorogenic detection of Serotonin (through the adduct serotonin-phhaldialdehyde) on DEC221 bilayer (excitation 488 nm, emission 500-650nm). (A) 100 µM ortho-phthalaldehyde on bilayer. (B) 100 µM phthalaldehyde added to pre-incubated serotonin (100 µM) on the bilayer. (C) Intensity analysis of serotonin OPA adduct on disordered and ordered domain. (D) is the AFM image of a DEC221 phase separated bilayer and (E) is the corresponding confocal image of the same area having fluorescence of serotonin-OPA adduct. (F) is the height profile of the AFM image.

**<u>Fig. S3</u>**

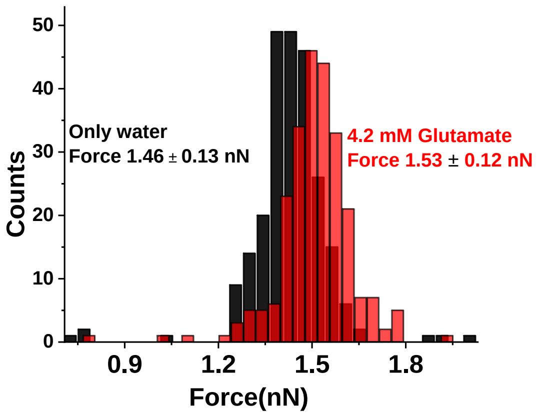


Fig. S3: Indentation force histogram of PPC111 bilayer in presence of water only (black), and of 4.2 mM of Glutamate (red)

**Fig. S4**

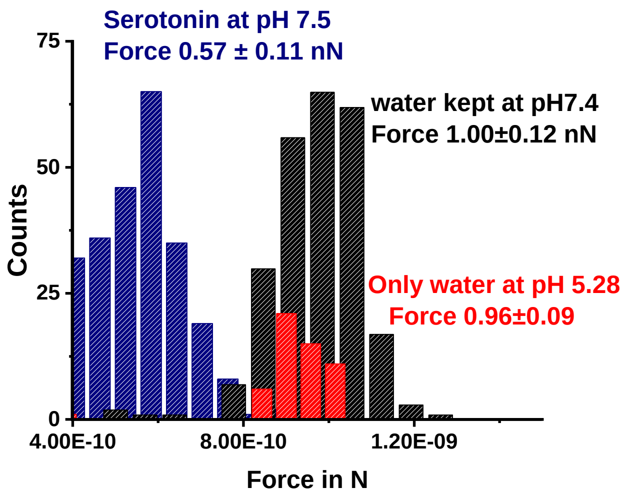


Fig. S4: Indentation force histogram of PPC111 bilayer in water at pH 5.28 (red), force histogram of the same bilayer in water after changing the pH to 7.4, before (black) and after (blue) addition of 4.1 mM of serotonin at pH 7.4.

**Fig. S5**

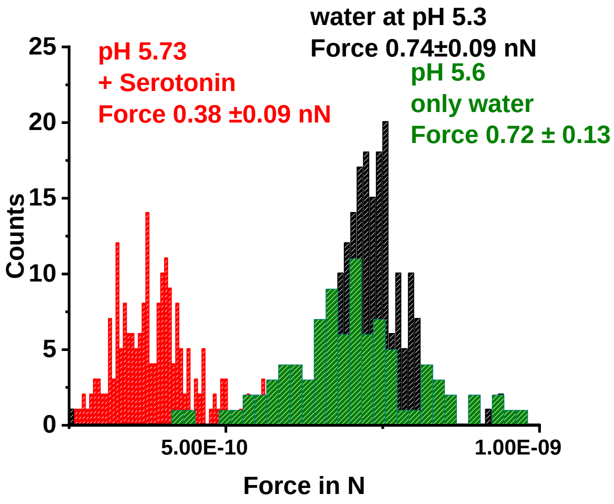


Fig. S5: Indentation force histogram of PPC111 bilayer in water at pH 5.3 (black), force histogram of the same bilayer in water having pH 5.6 (equivalent to serotonin pH); before (green) and after (red) addition of 4.1 mM of serotonin at pH 5.73.

**Fig. S6**

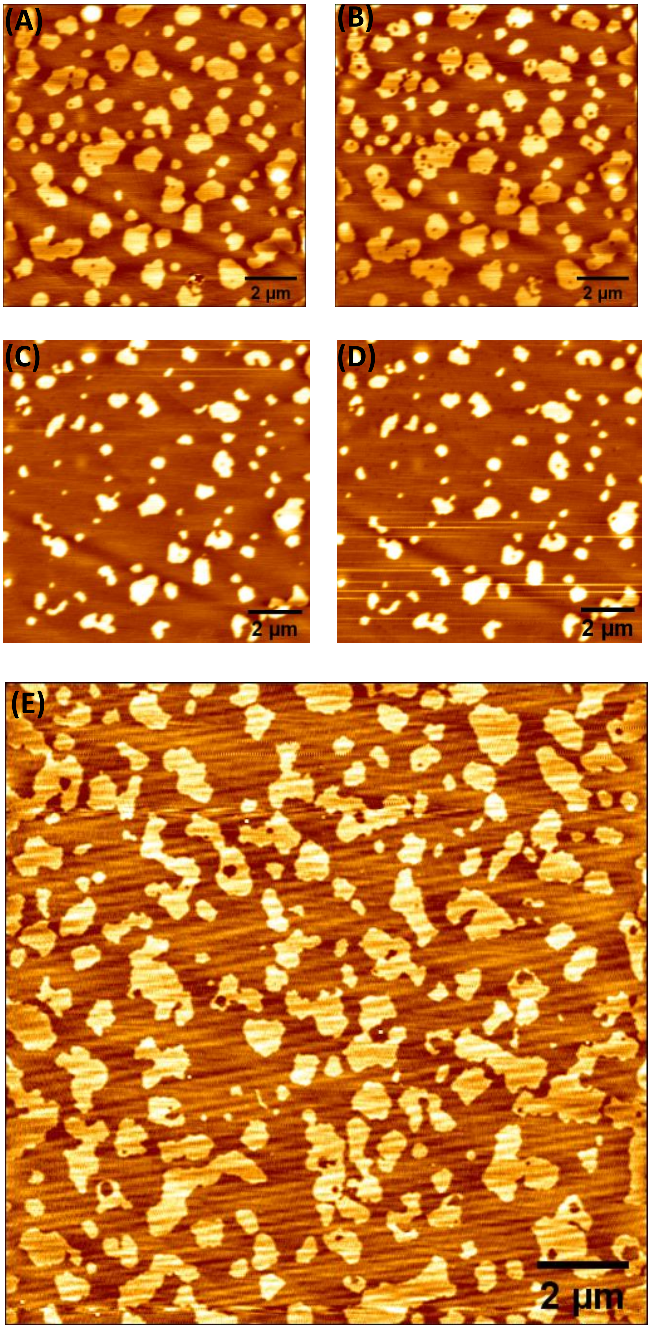


Fig. S6: AFM surface topography of DEC221 bilayer (A) without serotonin, (B) 2min, (C) 180 mins (D) 400 minutes after incubation of 4.1 mM serotonin. (E) A different bilayer prepared simultaneously kept for equivalent amount of time without serotonin treatment and imaged after 480 minutes.

**Fig. S7**

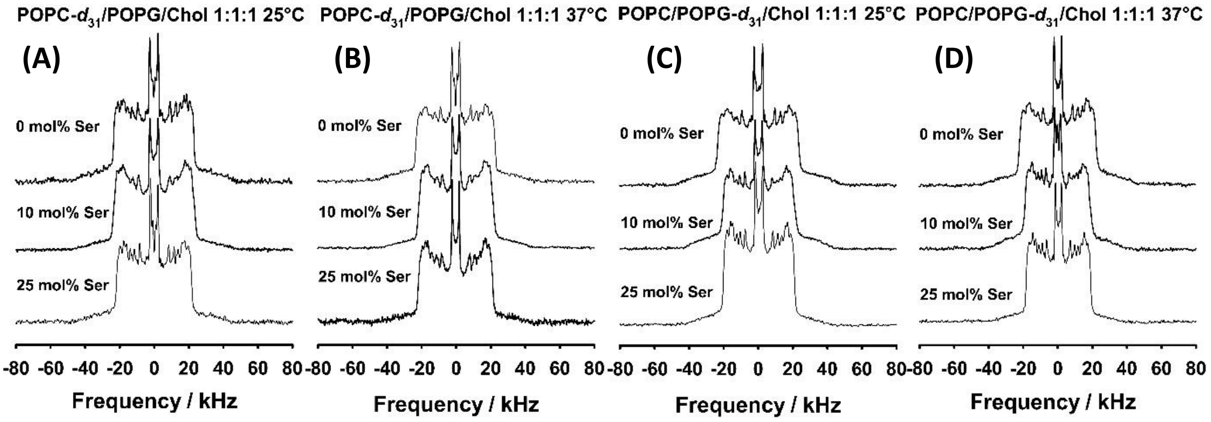


Fig. S7: (A) $^{2}$H NMR spectra of POPC-$d_{31}$:POPG:Chol(1:1:1)-Serotonin at (0, 10, 25 mol%) multilamellar vesicles at 25° C and (B) at 37° C, (C) $^{2}$H NMR spectra of POPC:POPG-$d_{31}$:Chol(1:1:1)-Serotonin multilamellar vesicles at (0,10, 25 mol%) at 25° and (D) at 37°.

### Fig. S8

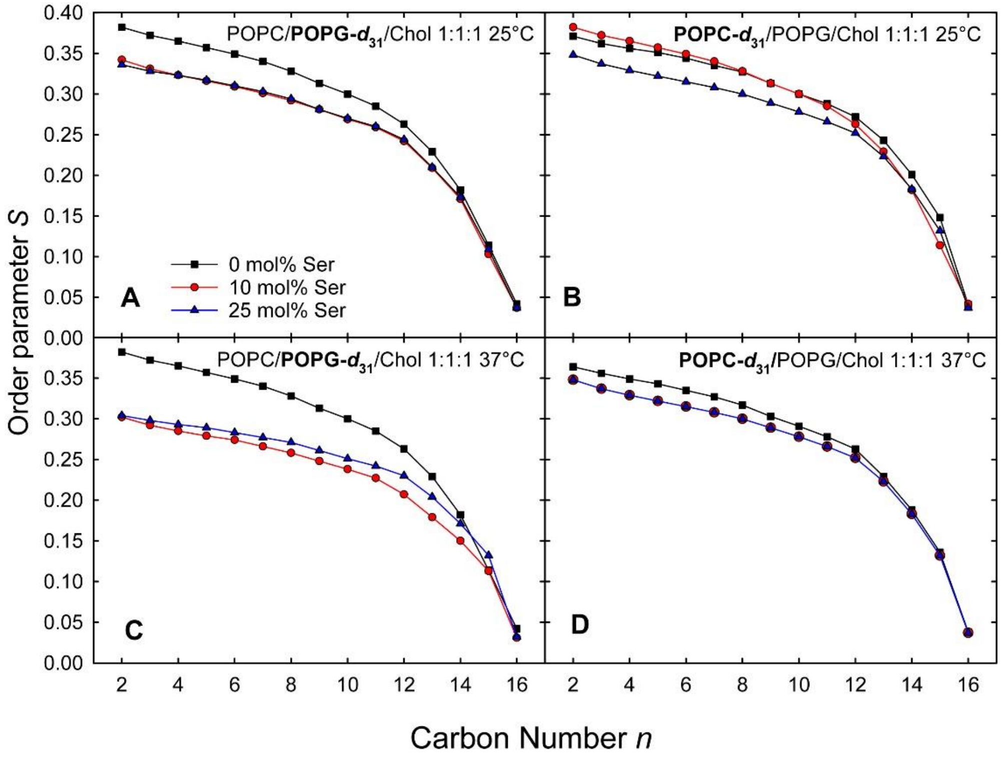


Fig. S8: $^2$H NMR order parameter profiles of POPC:POPG-$d_{31}$:Chol(1:1:1)-Serotonin at (0, 10, 25 mole percent at (A) 25°C and (C) 37°C. (B,D) are order parameter profiles of POPC-$d_{31}$:POPG:Chol(1:1:1)-Serotonin at 25°C and 37°C respectively.

**Fig. S9**

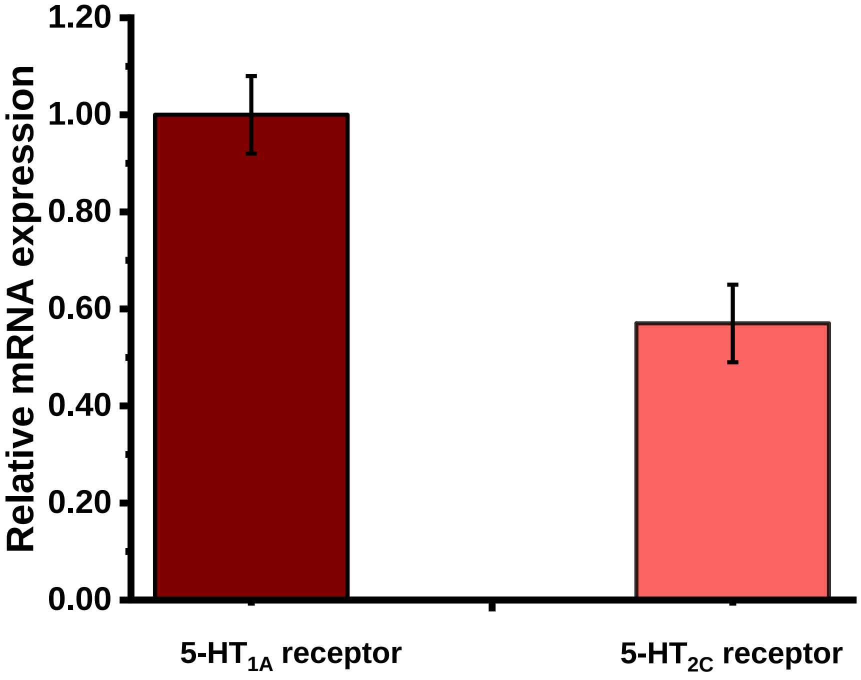


Fig. S9: Relative mRNA expression of 5-HT receptors in RN46A cells. Graph depicts quantitative qPCR analysis of mRNA expression of 5-$HT_{1A}$ and 5-$HT_{2C}$ receptors normalized to the endogenous 18S ribosomal RNA gene, computed by the $\Delta\Delta$Ct method. mRNA expression levels of other 5-HT receptors (5-HT1B, 5-HT3, 5-HT4 and 5-HT7 receptors) analyzed did not cross the experimental cycle threshold cut-off value, and levels were not significant. Data are represented as fold change ± SEM (Representative results from n=3. Experiment was repeated twice N=2).

**Fig. S10**

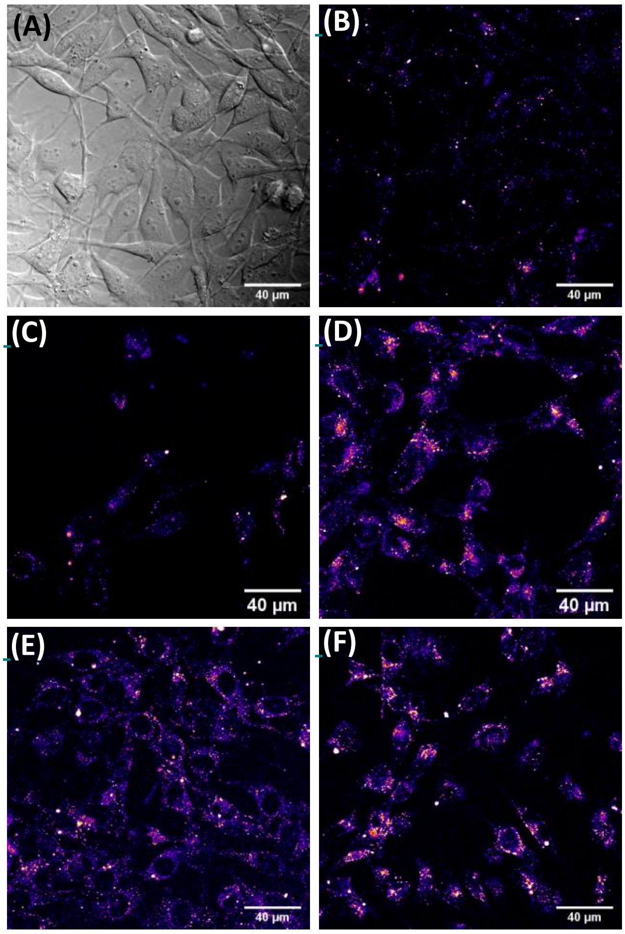


Fig. S10: (A-F) Serotonin-modulated binding of Rhodamine (Rh)- labelled IAPP oligomers to RN46A cells. (A) Transmission image of cells (B). (B) Auto fluorescence from the cells. (C) Cells were pre-treated with serotonin receptor blockers, 5-$HT_{1A}$ receptor antagonist WAY100635 (10µM), 5-$HT_2$ receptor antagonist Methysergide (10µM), 5-$HT_2$ receptor antagonist Ketanserin (10µM) and the 5-HT transporter (SERT) blocker, Fluoxetine (10µM); (D) cells incubated with only Rh-IAPP (200nM) for 30 minutes. (E)Cells pre-treated with serotonin receptor blockers + SERT blockers followed by Rh-IAPP, (F) Cells pre-treated with serotonin receptor blockers + SERT blockers + serotonin followed by Rh-IAPP.

**Fig. S11**

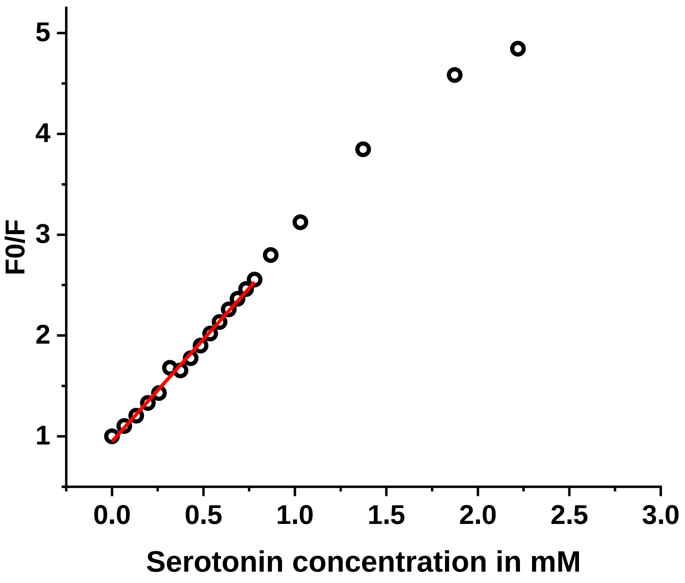


Fig. S11: Estimation of quenching of Rh-IAPP bound to PPC111 vesicles by serotonin. The Stern-Volmer plot is fitted (red line) upto ~1 mM serotonin concentration.
